# TDP-43 proteinopathy alters the ribosome association of multiple mRNAs including the glypican Dally-like protein (Dlp)/GPC6

**DOI:** 10.1101/2020.07.01.182360

**Authors:** Erik M Lehmkuhl, Suvithanandhini Loganathan, Eric Alsop, Alexander D Blythe, Tina Kovalik, Nicholas P Mortimore, Dianne Barrameda, Chuol Kueth, Randall J Eck, Bhavani B Siddegowda, Archi Joardar, Hannah Ball, Maria E Macias, Robert Bowser, Kendall Van Keuren-Jensen, Daniela C Zarnescu

**Affiliations:** Department of Cellular and Molecular Biology, University of Arizona, Tucson, AZ; Translational Genomics Research Institute, Phoenix, AZ; Department of Neurobiology, Barrow Neurological Institute, Phoenix, AZ; Department of Neuroscience, University of Arizona, Tucson, AZ; Department of Neurology, University of Arizona, Tucson, AZ

**Author notes:** **Corresponding Author:** Daniela C Zarnescu, PhD, Department of Cellular and Molecular Biology, University of Arizona, 1007 E. Lowell St, LSS RM 548A, Tucson, AZ 857 USA.

**Keywords:** ALS, TDP-43, translation, motor neuron, neuromuscular junction, glypican, Wnt signaling, Drosophila

## Abstract

Amyotrophic lateral sclerosis (ALS) is a genetically heterogeneous neurodegenerative disease in which 97% of patients exhibit cytoplasmic aggregates containing the RNA binding protein TDP-43. Using tagged ribosome affinity purifications in *Drosophila* models of TDP-43 proteinopathy, we identified TDP-43 dependent translational alterations in motor neurons impacting the spliceosome, pentose phosphate and oxidative phosphorylation pathways. A subset of the mRNAs with altered ribosome association are also enriched in TDP-43 complexes suggesting that they may be direct targets. Among these, *dlp* mRNA, which encodes the glypican Dally like protein (Dlp)/GPC6, a wingless (Wg/Wnt) signaling regulator is insolubilized both in flies and patient tissues with TDP-43 pathology. While Dlp/GPC6 forms puncta in the *Drosophila* neuropil and ALS spinal cords, it is reduced at the neuromuscular synapse in flies suggesting compartment specific effects of TDP-43 proteinopathy. These findings together with genetic interaction data show that Dlp/GPC6 is a novel, physiologically relevant target of TDP-43 proteinopathy.

## Introduction

Amyotrophic lateral sclerosis (ALS) is a neurodegenerative disorder characterized by the progressive loss of motor neuron function, culminating in death due to respiratory failure^1, 2^. Mutations in several dozen genes including SOD1^3^, TDP-43^4, 5^, and C9orf72^6, 7^ have been implicated in disease pathogenesis, however for >79% of patients the cause of disease remains unknown^8^, suggesting that yet-to-be-uncovered mechanisms significantly contribute to the majority of ALS cases. At the cellular level, several processes have been linked to ALS including neuroinflammatory response^9^, cellular metabolism^10, 11^, RNA processing, axonal transport and protein homeostasis^12^. A hallmark of ALS is the finding that >97% of patient spinal cord motor neurons exhibit cytoplasmic puncta containing the RNA binding protein TDP-43 (TAR DNA Binding Protein 43)^13, 14^. Together with reports that 2-3% of ALS patients harbor TDP-43 mutations^4, 5, 14, 15^, these findings suggest a key role for TDP-43 in ALS pathogenesis regardless of etiology.

TDP-43 is a nucleo-cytoplasmic shuttling, DNA/RNA binding protein which is predominantly localized to the nucleus where it regulates transcription^16^ and splicing^17, 18^. In the cytoplasm, TDP-43 regulates stress granule dynamics^19, 20, 21, 22^ as well as axonal and dendritic mRNA localization and translation^23, 24, 25^. In disease, TDP-43 is depleted from the nucleus, causing splicing defects, derepression of cryptic exons^26^ and increased retrotransposon expression^27, 28, 29, 30^. TDP-43 has been shown to induce toxicity through both nuclear loss-of-function, and cytoplasmic gain-of-function, mechanisms^31, 32, 33^. When mislocalized to the cytoplasm, TDP-43 associates with a plethora of RNA containing complexes including stress and transport granules^34, 35, 36^, as well as protein complexes devoid of RNA^37, 38^. TDP-43 has also been shown to influence the translation of specific mRNAs, <u>as both a</u> negative and a positive regulator^24, 25, 39, 40, 41^. Taken together these findings suggest a complex role for TDP-43 in translation regulation, which has been linked to the formation of TDP-43 cytoplasmic puncta in disease and altered protein expression and/or localization of its mRNA targets.

A potential mechanism by which TDP-43 cytoplasmic inclusions dysregulate translation is by altering the ribosomes’ access to mRNAs, as suggested by the “ribostasis hypothesis”^42^. Further substantiating this model are recent reports of TDP-43 mediated translation inhibition of mRNAs, including *futsch* and *hsc70-4* mRNAs (in axons)^24, 39^ and *Rac1* mRNA (in dendrites)^25^ all of which were identified through candidate approaches. However, the relationship between TDP-43 associated mRNAs and translation in disease has not yet been examined through an unbiased approach. Here we combined RNA immunoprecipitations (RIP) and translating ribosome affinity purification (TRAP) to identify mRNAs that simultaneously satisfy two criteria: 1) are enriched in TDP-43 complexes and thus have the potential to be regulated by TDP-43, and 2) their association with ribosomes is altered in the context of *Drosophila* models of TDP-43 proteinopathy, consistent with an effect on translation. RIP experiments uncovered candidate mRNA targets linked to neuromuscular junction development and synaptic growth. TRAP uncovered mitochondrial metabolism, proteostasis and wingless signaling as significant components of the “normal” motor neuron translatome. When compared to controls, TDP-43 proteinopathy was found to significantly alter the ribosome association of multiple mRNAs whose cognate proteins play roles in splicing, purine metabolism and mitochondrial electron transport among others.

Here we identify Dally-like protein (Dlp), a glypican-type heparan sulfate proteoglycan (HSPG) that has been described as a regulator of wingless (Wg/Wnt)^43^ and Liprin alpha receptor (LAR) mediated signaling at the neuromuscular junction^44^ as a candidate target of TDP-43 mediated translation inhibition. We show that *dlp* mRNA is enriched in TDP-43 complexes and sequestered in insoluble aggregates, consistent with the ribostasis hypothesis. In line with this, we find that Dlp protein is significantly reduced at the neuromuscular junction while steady state levels of *dlp* transcript remain unchanged, further supporting the possibility of translation inhibition in axons and/or at synapses. Surprisingly, Dlp also accumulates in puncta within the ventral nerve cord neuropil, suggesting that TDP-43 proteinopathy affects Dlp in a compartment specific manner. Dlp depletion at the NMJ, but not puncta formation, was also observed with endogenous TDP-43 knockdown suggesting that some Dlp phenotypes are the result of loss of nuclear TDP-43 function while others are the results of toxic cytoplasmic gain of function. Genetic interaction experiments show that *dlp* overexpression mitigates TDP-43 dependent locomotor deficits, consistent with the notion that Dlp mediates aspects of TDP-43 proteinopathy *in vivo*. Lastly, GPC6 protein, a human homolog of Dlp, exhibits aggregate-like accumulations while GPC6 mRNA is enriched in insoluble fractions derived from ALS patient spinal cords mirroring the findings from Drosophila motor neurons. Together, these findings highlight key pathways altered in the motor neuron translatome in the context of TDP-43 proteinopathy and support the notion that altered expression of the glypican Dlp/GPC6 contributes to motor neuron degeneration.

## Results

### mRNAs enriched with TDP-43 in motor neurons encode proteins linked to neuronal and synaptic signaling

TDP-43 has been shown to associate with a plethora of RNA targets and regulate various aspects of RNA processing including RNA localization and translation^23, 24, 41^. These findings suggest that cytoplasmic TDP-43, which is a hallmark of disease, has multiple opportunities to cause alterations in the motor neuron proteome and contribute to pathogenesis. To uncover these alterations, we defined TDP-43 dependent changes in the motor neuron translatome *in vivo* using *Drosophila* models of TDP-43 proteinopathy that recapitulate key aspects of the disease including locomotor defects, cytoplasmic aggregates and reduced lifespan^45, 46^. We hypothesized that while some of the translational alterations may be an indirect consequence of neurodegeneration, others are directly caused by TDP-43, possibly through mRNA association with, TDP-43 cytoplasmic complexes <u>and consequent sequestration</u>. To distinguish between direct versus indirect effects on the translatome we set out to identify mRNAs that are both enriched with TDP-43 and translationally dysregulated in motor neurons *in vivo* using a combination of RNA immunoprecipitations (RIP) and Translating Ribosomes Affinity Purifications (TRAP)^47, 48^ in the context of TDP-43 proteinopathy. To this end, we first used *Drosophila* larvae expressing human TDP-43 protein specifically in motor neurons via the GAL4-UAS system (D42 GAL4 > UAS-TDP-43-YFP) and performed immunoprecipitation experiments to pull down TDP-43 and associated mRNAs (RIP). The mRNAs associated with TDP-43 (WT or mutant G298S) were isolated and subjected to RNA sequencing (see Materials and Methods, and Figure 1a). Bioinformatic analyses identified several mRNAs significantly enriched with TDP-43, as determined by comparing TDP-43 associated mRNAs in a complex with the transcriptome of dissected ventral nerve cords expressing the appropriate TDP-43 variant (TDP-43^WT^ ^or^ ^G298S^, see Figures 1b,c). Of the approximately 9,900 transcripts detected in ventral nerve cords (Supplemental Table 1-1), 1,892 were enriched with, and shared by, both TDP-43 variants (Log2FC>2, Padj <0.05). Additionally, about 25% of mRNAs enriched with TDP-43 were unique to TDP-43^WT^ or TDP-43^G298S^ (662 mRNAs enriched with TDP-43^WT^ and 616 mRNAs enriched with TDP-43^G298S^, respectively, see Figure 1d) consistent with previous evidence that wild-type and mutant TDP-43 variants cause toxicity in part through distinct mechanisms (see also Figure 1e-f)^39^.

**Figure 1.**
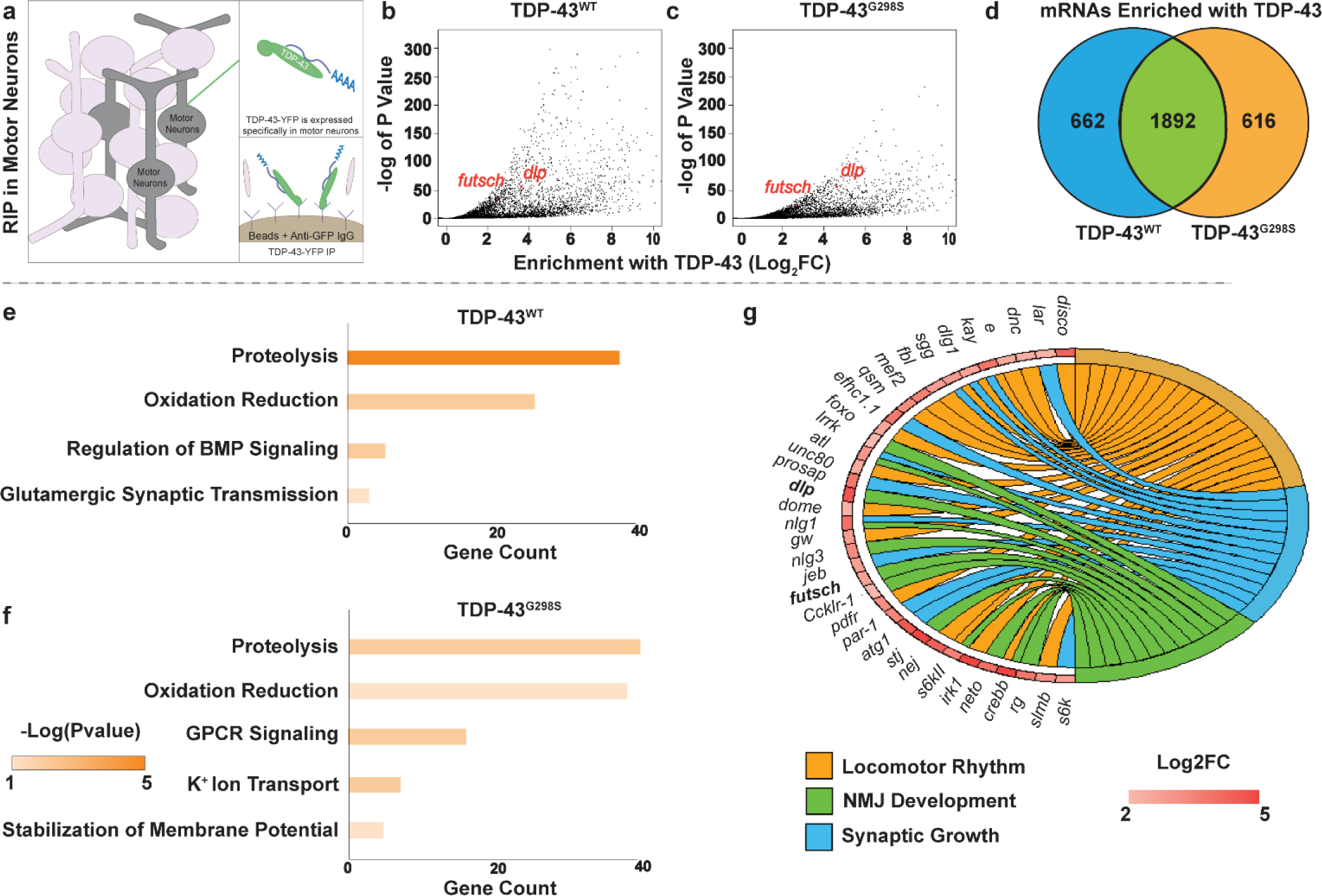
mRNAs Enriched with TDP-43 in A Drosophila Model of ALS. **(a)**. Experimental schematic for RNA immunoprecipitations of human TDP-43, specifically from the motor neurons of third instar larvae. **(b**,**c)**. Volcano plot displaying mRNAs enriched with TDP-43 in TDP-43^WT^ **(b)** and TDP-43^G298S^. All genes associated with TDP-43 (log2FC > 0) are displayed regardless of significance. **(c)** proteinopathy, relative to transcript levels in the ventral nerve cord. **(d). (e**,**f**,**g)**. GO terms for TDP-43 enriched genes (log2FC > 2, Padj <0.05) either unique to TDP-43^WT^ **(e)** or TDP-43^G298S^ **(f)** or shared between genotypes **(g)**.

GO term analyses using David 6.8^49, 50^ of mRNAs enriched with both TDP-43^WT^ and TDP-43^G298S^ (Log2FC>2, Padj <0.05) highlight neuronal pathways previously linked to TDP-43 such as NMJ development^51^ and synaptic growth^52, 46^. (Figure 1e-g, Supplemental Table 1-2,3,4) and are consistent with published studies of TDP-43 interacting RNAs in a murine model of TDP-43 proteinopathy^53^. In addition, our analyses identified previously published mRNA targets of TDP-43 including *futsch*^24^ (TDP-43^WT^, Log2FC = 2.44, Padj = 8.07E-37; TDP-43^G298S^, Log2FC = 2.68, Padj = 6.86E-12) and *CSNK1E*^54^ (TDP-43^WT^, Log2FC = 0.721, Padj = 3.32E-3; TDP-43^G298S^, Log2FC = 0.847, Padj = 3.08E-3). Furthermore, we found that of the 834 TDP-43 interacting mRNAs recently identified in murine neurons using TRIBE (targets of RNA-binding proteins identified by editing)^55^ about 1/3 were also enriched in our *Drosophila* RIP data sets (Supplemental Table 1-5). In summary, our RIP experiments identify several mRNA candidate targets of TDP-43 in motor neurons, *in vivo*, a fraction of which have been previously identified in mammalian neurons^55, 24, 54^ while others are novel, and together they highlight links between TDP-43 proteinopathy and neuronal function, neuromuscular junction development and synaptic growth.

### TDP-43 proteinopathy induces a broad range of translational changes in motor neurons

Next, to define the *in vivo* motor neuron translatome and identify changes in translation induced by TDP-43 proteinopathy (see Materials and Methods, and Figure 2a) we conducted TRAP experiments^47, 48^. To this end, RpL10-GFP was expressed in motor neurons using D42 GAL4 either on its own, or together with TDP-43^WT^, or TDP-43^G298S^. Following ribosome immunoprecipitations we isolated total RNA and conducted RNA seq, detecting ∼9,500 mRNAs (9,711 mRNAs for RpL10 controls, 9,347 for TDP-43^WT^ and 9,669 for TDP-43^G298S^, see Materials and Methods, Supplemental Table 2-1). We first defined the “normal” motor neuron translatome using the normalized gene counts associated with ribosome immunoprecipitations from RpL10-GFP controls (Supplemental Table 2-2). Using David 6.8^49, 50^, we determined that genes enriched in the top 10% of the motor neuron translatome normalized counts were related to proteostasis and energy metabolism (Figure 2b, Supplemental Table 2-3), consistent with high levels of protein turnover^56^ and energy demands in neurons^57^.

**Figure 2.**
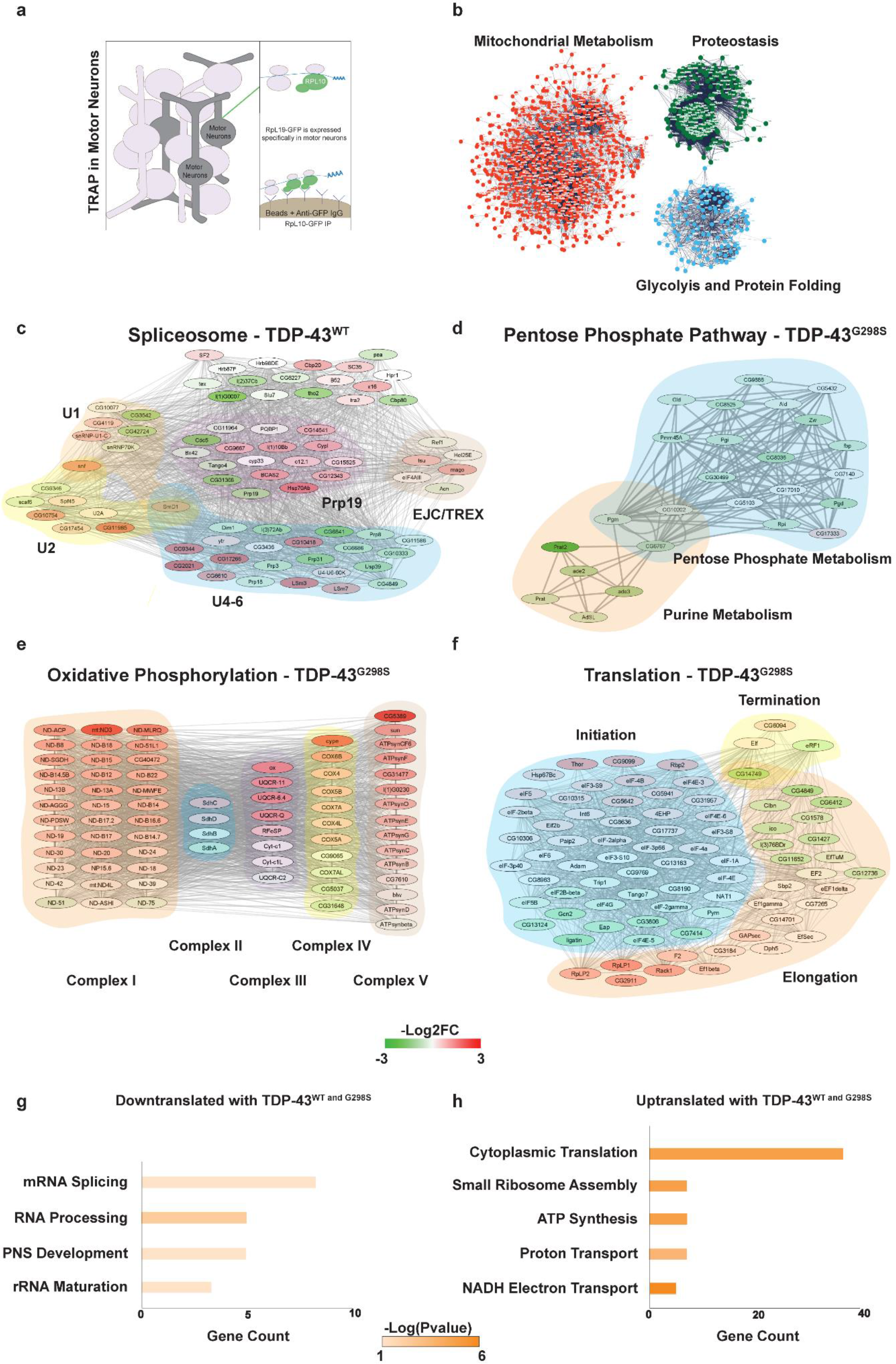
Translational Alterations Induced by TDP-43 Proteinopathy. **(a)**. Experimental schematic for RNA immunoprecipitations of human TDP-43, specifically from the motor neurons of third instar larvae. **(b)**. STRING clusters from genes in the top 10% of normalized counts associated with RpL10-GFP in w^1118^ larvae (control motor neuron translatomes). **(c)**. Altered ribosome association of spliceosome genes in TDP-43^WT^ relative to the control. **(d)**. Altered ribosome association of purine metabolism genes in TDP-43^G298S^ relative to control. **(e)**. Altered ribosome association of oxidative phosphorylation genes in TDP-43^G298S^ relative to control. **(f)**. Altered ribosome association of translation associated genes in TDP-43^G298S^ relative to control. **(g**,**h)**. GO terms for RpL10 depleted genes (log2FC < −1) **(g)** and RpL10 enriched genes (log2FC >1) **(h)** shared between both TDP-43^WT^ and TDP-43^G298S^ models.

To identify changes in translation caused by TDP-43 proteinopathy in motor neurons *in vivo*, we subsequently compared the mRNAs enriched with ribosomes precipitated from *Drosophila* larvae expressing RpL10-GFP TDP-43^WT^ or RpL10-GFP TDP-43^G298S^ to RpL10-GFP controls after normalizing to the transcriptomes of dissected ventral nerve cords of the appropriate genotype (RpL10-GFP TDP-43^WT^, RpL10 GFP TDP-43^G298S^ or RpL10-GFP). These comparisons identified several genes enriched with or depleted from motor neuron ribosomes in both of the ALS models relative to RpL10-GFP controls (ΔLog2FC >1 or <-1),. Subsequent GO term analyses using David 6.8 ^49, 50^ revealed that in the context of TDP-43^WT^, spliceosome components were translationally dysregulated (Figure 2c, Supplemental Table 2-4), suggesting that TDP-43^WT^ overexpression causes pathway alterations previously associated with nuclear loss -of-function phenotypes^58, 59^. Interestingly, in the context of TDP-43^G298S^, we found significant alterations in neuronal metabolism (Supplemental Table 2-5), including purine processing leading into the pentose phosphate pathway (PPP), which provides a mechanism for countering oxidative stress via increased NADPH production^60^. PPP was found to down-translated, consistent with our previous studies showing that glucose 6 phosphate dehydrogenase, the rate limiting enzyme in PPP is altered in flies and patient tissues with TDP-43 pathology^61^ (Figure 2d). Interestingly, oxidative phosphorylation genes encoding components of the electron transport chain showed both increased and reduced association with ribosomes (Figure 2e). Recent findings of reduced Complex I activity would suggest that uptranslation of Complex I components identified via TRAP may reflect a compensatory mechanism^10, 11, 62, 63^.

Interestingly, genes salient to translation were also dystranslated (Figure 2f). In both the TDP-43^WT^ and TDP-43^G298S^ models, ribosomal RNA maturation was down-regulated (Figure 2g, Supplemental Table 2-6) while cytoplasmic translation components were up-regulated relative to the control (Figure 2h, Supplemental Table 2-7). These findings are consistent with a previous study in which patient derived iPSC neurons exhibited a compensatory increase in global translation^64^. Further corroborating our findings with previous studies, we detected increased association with ribosomes for *CG6762*, the *Drosophila* ortholog of human SRXN1, (TDP-43^WT^, Log2FC = 0.673, Padj = 2.79E-32; TDP-43^G298S^, Log2FC = 0.328, Padj = 2.50E-51), one of 14 genes that exhibited increased translation in a human cell model of TDP-43^A315T^ proteinopathy^41^. Taken together, our ribosomal tagging experiments highlight complex changes in translation, some of which may be direct consequences of TDP-43 proteinopathy while others may reflect compensatory mechanisms.

### A fraction of TDP-43 associated mRNAs are altered in their association with ribosomes in the context of TDP-43 proteinopathy

To distinguish between translational changes that may be caused directly by TDP-43 versus compensatory alterations, we compared genes altered in their association with ribosomes to those enriched in TDP-43 complexes, in the context of TDP-43 proteinopathy. These analyses that approximately 1/3 of the genes depleted from ribosomes were also enriched in TDP-43 complexes (Figure 3a, Supplemental Table 3-1, TDP-43^WT^ 30.6%, TDP-43^G298S^ 37.6%) suggesting that these may be direct targets of translation inhibition. GO term analyses of these TDP-43 enriched, TRAP depleted genes using David 6.8^49, 50^ identified pathways that have previously been associated with ALS including GPCR signaling^65, 66^, NMJ Development^65, 66^, Autophagy^67, 68^, ER Organization^69^, Immune Respone^70^ and Oxidation Reduction^71^ among others (Figure 3b-e, Supplemental Table 3-2,3). Interestingly, a similar proportion (Figure 3f, Supplemental Table 3-4, TDP-43^WT^ 27.6%, TDP-43^G298S^ 34.9%) of the genes enriched with ribosomes in the context of TDP-43 proteinopathy were also associated with TDP-43 complexes (Figure 3f, Supplemental Table 3-4), consistent with previous finding that TDP-43 can also function as a positive regulator of translation^41^. GO term analyses identified pathways such as Membrane Potential Stability and Transmembrane Transport – K+ (Figure 3g-j, Supplemental Table 3-4) comprising genes related to membrane excitability, a process known to be altered in ALS^72^. We note that although GO term analysis is useful to identify translational alterations affecting multiple components of the same pathway, the genes encompassed by these GO terms represent only a fraction of genes enriched and translationally dysregulated in the context of TDP-43 (Figure 3a, f, TDP-43^WT^ = 13.7%, TDP-43^G298S^ = 12.4%). This suggests that in addition to specific pathways, a plethora of individual genes might represent salient targets of TDP-43 mediated translational inhibition or activation.

**Figure 3.**
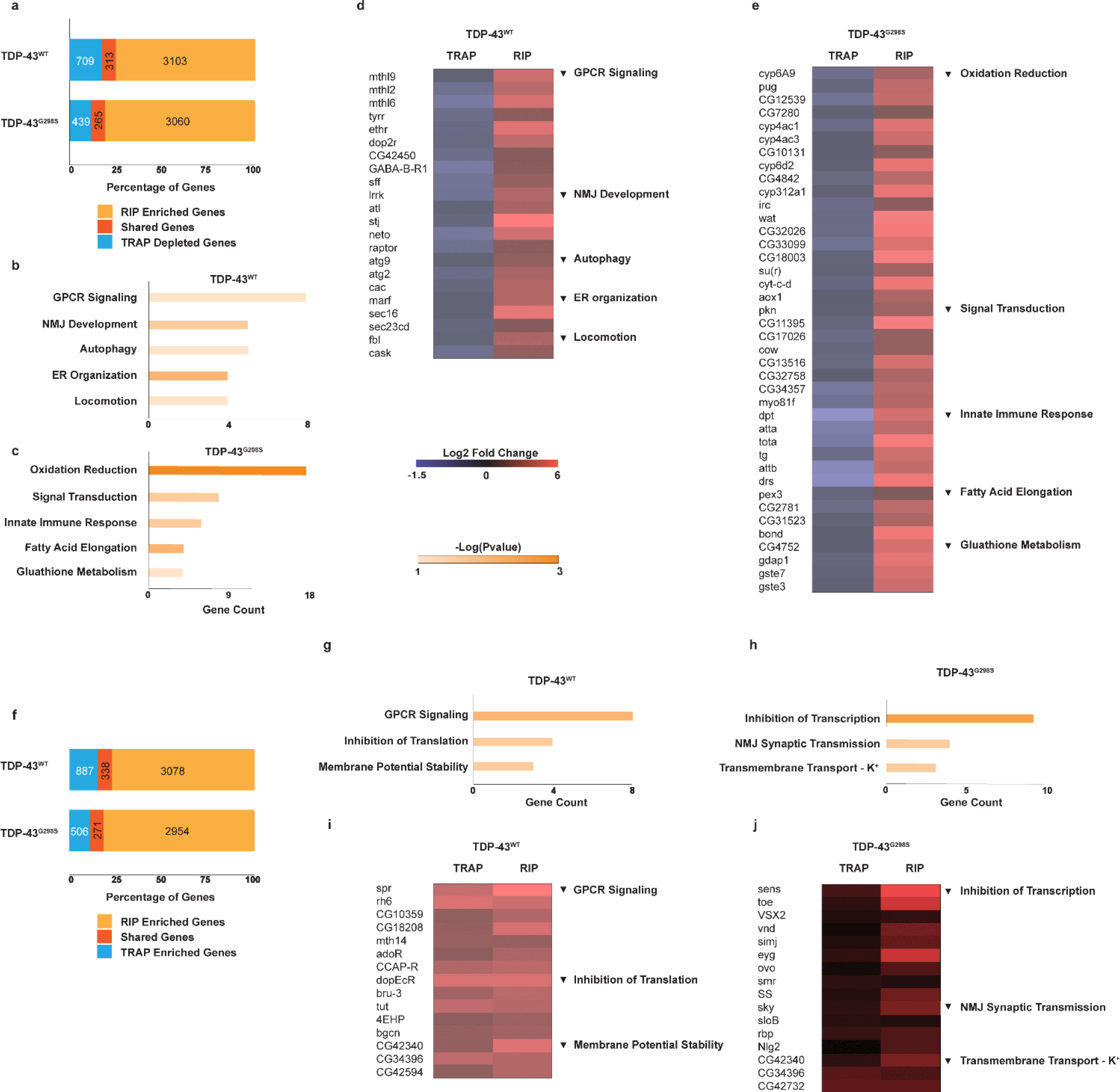
Direct Targets of TDP-43 Mediated Translation Regulation. **(a)**. Overlap between genes enriched with TDP-43 (log2FC >1, padj <0.05) and depleted with RpL10 (log2FC <-1) relative to the w^1118^ control. **(b**,**c)**. GO terms for TDP-43 enriched, RpL10 depleted genes.in TDP-43^WT^ **(b)** or TDP-43^G298S^ **(c)** models. **(d**,**e)**. RpL10 depletion (TRAP) and TDP-43 enrichment (RIP) for genes constituting significant GO terms in TDP-43^WT^ **(d)** and TDP-43^G298S^ **(e). (f)**. Overlap between genes enriched with TDP-43 (log2FC >1, padj <0.05) and enriched with RpL10 (log2FC > 1) relative to the w^1118^ control. **(g**,**h)**. GO terms for TDP-43 enriched, RpL10 enriched genes.in TDP-43^WT^ **(g)** or TDP-43^G298S^ **(h)** models. **(i**,**j)**. RpL10 enrichment (TRAP) and TDP-43 enrichment (RIP) for genes constituting significant GO terms in TDP-43^WT^ **(i)** and TDP-43^G298S^ **(j)**.

### *dally-like protein* (*dlp*) mRNA, a glypican involved in wingless (Wg/Wnt) signaling is sequestered in insoluble complexes in *Drosophila* models of TDP-43 proteinopathy

Additional analyses of the “normal” motor neuron translatome using STRING^73^ identified an enrichment in Wg/Wnt signaling genes (Figure 4a), with the top 5% of genes translated in motor neurons containing Wg/Wnt signaling genes at a rate 61% greater than that of the entire data set. This is consistent with the established role of Wg/Wnt signaling in motor neurons and at the neuromuscular junction^74^. Interestingly, Wg/Wnt signaling has been shown to be dysregulated in ALS^75, 76, 77^, although its mechanisms remain poorly understood, in part due to the complexity of Wg/Wnt signaling and cross-talk with other pathways. Given these observations, we queried the pathway for candidate targets of TDP-43 mediated translational inhibition and found *dlp* mRNA to be enriched in TDP-43 complexes (TDP-43^WT^, Log2FC = 3.62, Padj = 8.32E-57; TDP-43^G298S^, Log2FC = 4.63, Padj = 2.47E-57) and significantly depleted from ribosomes in the context of TDP-43 proteinopathy (TDP-43^WT^, Log2FC = −1.03, Padj = 0.027; TDP-43^G298S^, Log2FC = −0.85, Padj = 0.012, see also Supplemental Table 4-1). *dlp* encodes a glypican, a member of the heparan sulfate proteoglycan (HSPG) family, localized to the plasma membrane via a GPI anchor or secreted in the extracellular matrix. HSPGs consist of a protein core and several heparan sulfate (HS) glycosaminoglycan (GAG) linear polysaccharide chains which exhibit complex sulfation patterns that dictate the ability of the HSPGs to interact with ligands such as *wingless*^43^. In *Drosophila*, Dlp was shown to function as a co-factor and competitive inhibitor for binding to the Wg/Wnt pathway receptor Frizzled 2^43^. Interestingly, additional components of Wg/Wnt signaling exhibit altered ribosomal association including *frizzled 2* mRNA (TDP-43^WT^, Log2FC = −1.00, Padj. = 0.026; TDP-43^G298S^, Log2FC = −0.56, Padj. = 0.027) and *ovo/shavenbaby* mRNA (TDP-43^WT^, Log2FC = 2.74, Padj. = 4.38E-8; TDP-43^G298S^, Log2FC = 3.02, Padj. = 2.15E-21) (Figure 4b), further substantiating the possibility that TDP-43 proteinopathy affects this signaling pathway.

**Figure 4.**
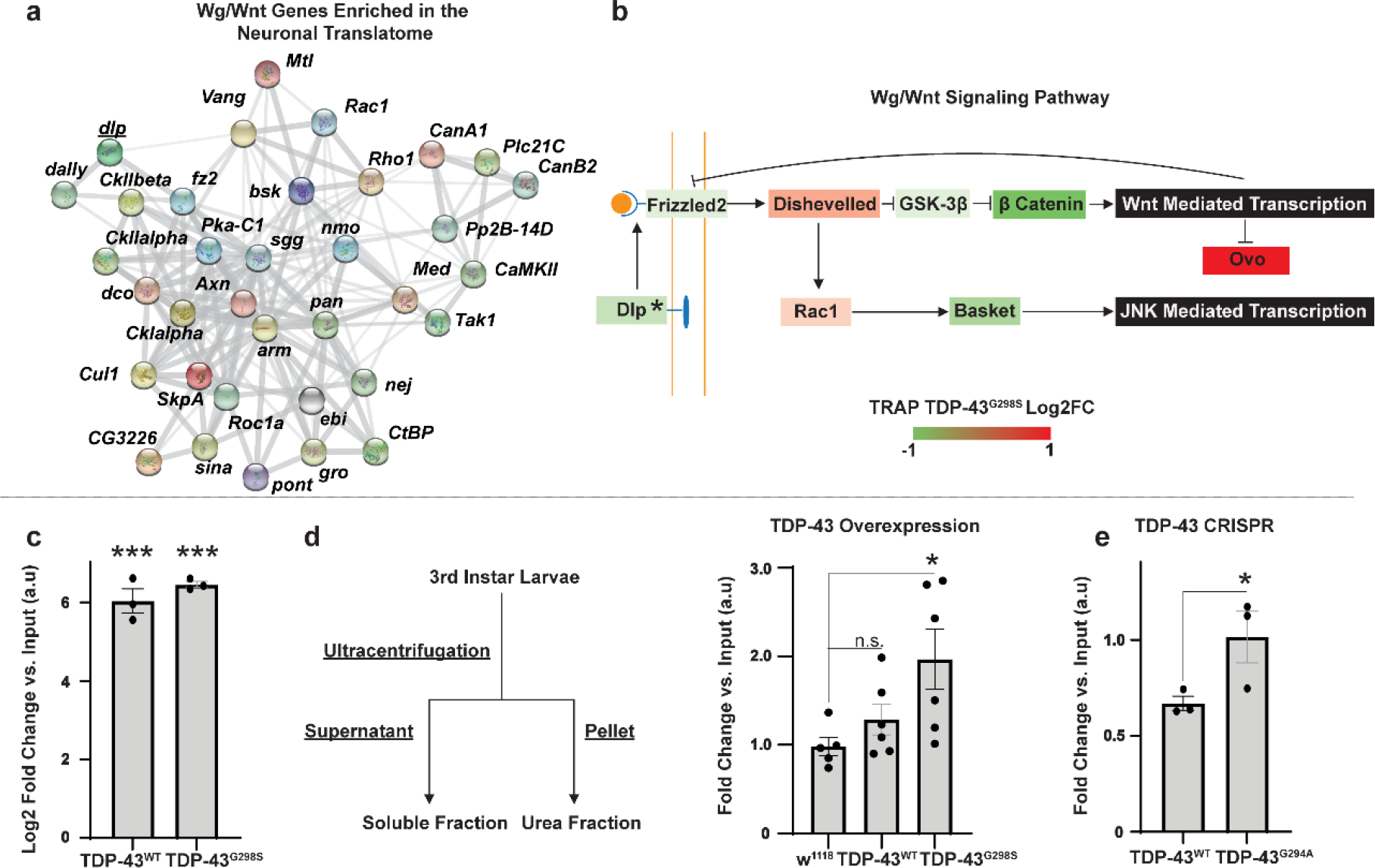
*dlp* mRNA, a Wg/Wnt interactor, is enriched in TDP-43 complexes and sequestered in insoluble fractions. **(a)**. STRING diagram ^73^ depicting relationship between Wg/Wnt signaling genes in the top third of the TRAP control IP normalized counts (D42>RpL10 GFP, see Materials and Methods). Line thickness between genes is correlated with the strength of the interaction as defined by STRING. **(b)**. Altered ribosome association of a subset of Wnt signaling genes in TDP-43^G298S^ relative to control. **(c)**. qPCR quantification of *dlp* mRNA enrichment with TDP-43 complexes in TDP-43^WT^ and TDP-43^G298S^ expressing larvae (D42>TDP-43 WT or G298S). Enrichment in TDP-43 complexes is shown as arbitrary units (*a*.*u*.) after normalization to input. **(d**,**e)**. Schematic of fractionation experiments and qPCR quantification of *dlp* mRNA levels in the insoluble fraction (Urea fraction) of TDP-43^WT^ or TDP-43^G298S^ expressing larvae relative to w^1118^ controls. N = (w^1118^ = 5, other = 6) **(d)**, and CRISPR-TDP-43^G294A^ (relative to CRISPR-TDP-43^WT^, N = 3). Significance was determined using the Holm Sidak’s multiple comparisons test **(b**,**d)** or Student’s T Test. **(e)** (t_value_ = 2.480, degrees of freedom = 4). * = P_value_ < 0.05, ** = P_value_ < 0.01, *** = P_value_ < 0.001. Error bars represent SEM.

To validate *dlp* mRNA as a target of TDP-43 in motor neurons, we first performed TDP-43 immunoprecipitations from *Drosophila* larvae expressing either TDP-43^WT^ or TDP-43^G298S^ in motor neurons then isolated the associated RNA and used RT-qPCR to amplify the *dlp* transcript. These experiments showed that indeed, *dlp* mRNA was enriched with both TDP-43^WT^ (Log2FC = 6.11, P_value_ = 5.54E-7) and TDP-43^G298S^ (Log2FC = 6.46, P_value_ = 5.54E-7) relative to input (Figure 4c), confirming the RIP mRNA seq data (Figure 1a-d).

Next, to determine whether *dlp* mRNA is insolubilized as predicted by the ribostasis hypothesis, we conducted subcellular fractionations and quantified *dlp* mRNA in the soluble and insoluble/urea fractions from third instar larvae expressing TDP-43 in motor neurons. These experiments showed that *dlp* mRNA is significantly insolubilized in the context of TDP-43^G298S^ (Fold change = 1.99, P_value_ = 0.0256, Figure 4d) but not in the context of TDP-43^WT^, suggesting that although *dlp* associates with TDP-43 complexes regardless of genotype, the extent of insolubilization is variant dependent. Further substantiating this finding, *dlp* mRNA was insolubilized in a TDP-43^G294A^ (Fold change = 2.51, P_value_ = 0.034, Figure 4e) but not in the TDP-43^WT^ CRISPR fly model of ALS in which the endogenous *Drosophila* TBPH gene has been replaced with human TDP-43^78^. Taken together these results indicate that *dlp* mRNA is a novel TDP-43 candidate target that associates with insoluble aggregates in the context of TDP-43 proteinopathy and are consistent with previous reports of Wg/Wnt alterations in disease.

### Dlp protein levels are altered in the neuromuscular system in the context of TDP-43 proteinopathy

A possible outcome of *dlp* mRNA enrichment with TDP-43 complexes, insolubilization and decreased association with ribosomes is inhibition of Dlp protein synthesis. To test the impact of TDP-43 proteinopathy on Dlp protein expression we conducted immunofluorescence experiments in the larval neuromuscular system. We examined Dlp expression at the neuromuscular junction (NMJ) and the ventral nerve cord (VNC), which comprises motor neuron cell bodies and compactly packaged neurites, particularly dendrites^79^ (Figure 5a-z). These experiments showed that overexpression of TDP-43^WT^ or TDP-43^G298S^, and RNAi knock-down of TBPH (endogenous *Drosophila* TDP-43, TBPH^RNAi^) are sufficient to deplete Dlp protein within NMJ boutons relative to the w^1118^ control (TDP-43^WT^, decreased 35.7%, P_value_ = 0.00920; TDP-43^G298S^, decreased 37.2%, P_value_ = 0.00860; TBPH^RNAi^, decreased 42.5%, P_value_ = 0.00920; Figure 5m). While no major changes in Dlp expression were detected in the motor neuron cytoplasm within the VNC (data not shown), striking aggregate-like structures were observed in the neuropil when either TDP-43^WT^ or TDP-43^G298S^ were overexpressed (see Figures 5n-y). Quantification of Dlp granularity shows that both TDP-43^WT^ and TDP-43^G298S^ proteinopathy cause a statistically significant increase in Dlp granule number (TDP-43^WT^, Fold Change = 4.99, P_value_ = 1.77E-4; TDP-43^G298S^, Fold Change = 4.24, P_value_ = 8.49E-4) and cumulative granule area relative to the w^1118^ controls. In contrast, TBPH^RNAi^ did not cause significant Dlp granularity phenotypes (TBPH^RNAI^, Fold Change = 2.03, P_value_ = 0.262, Figures 5n-z and Supplemental Figure 5-1a).

**Figure 5.**
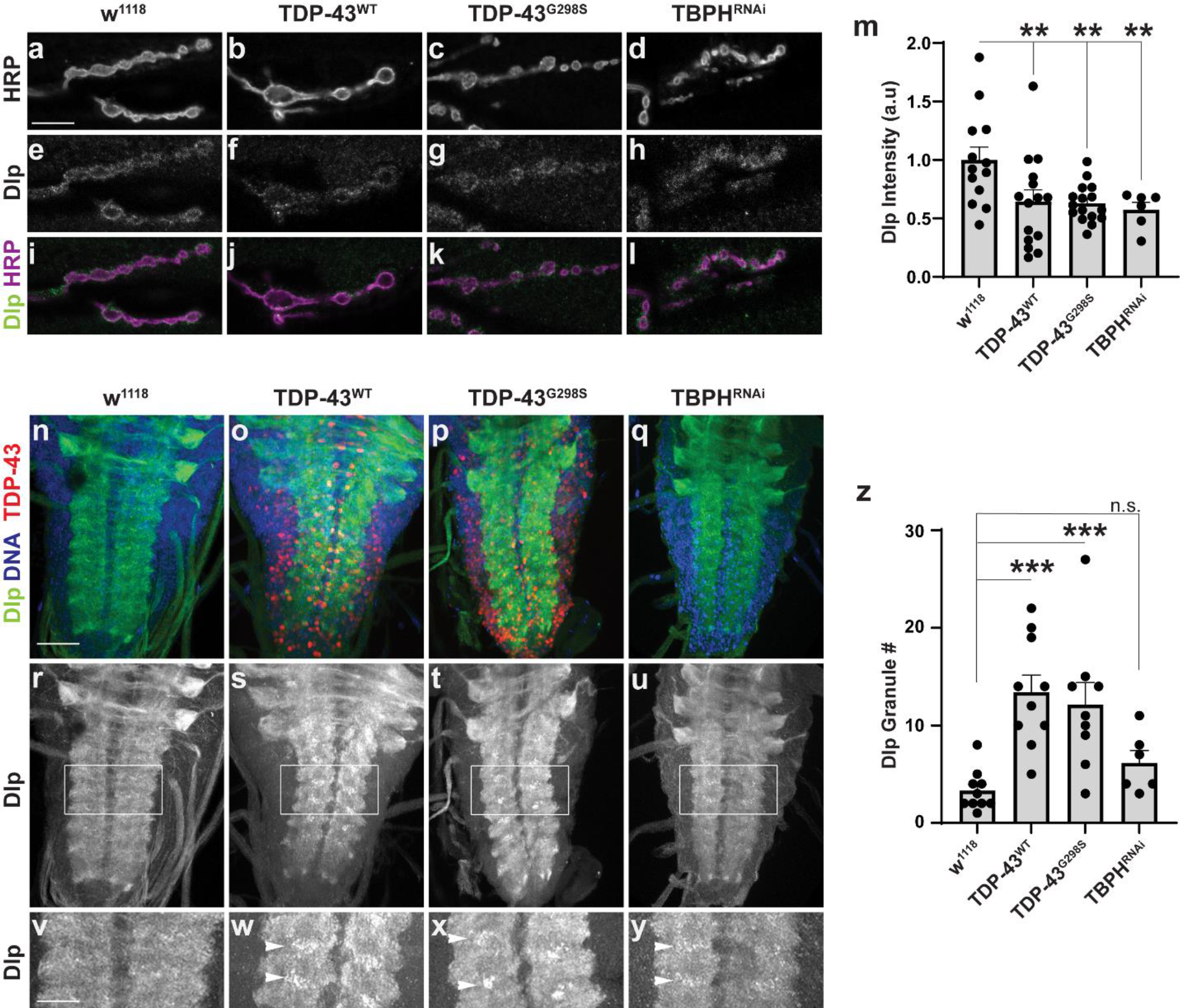
Dlp protein expression is altered by TDP-43 proteinopathy. **(a-l)** Representative images of w^1118^, TDP-43^WT^, TDP-43^G298S^ and TBPH^RNAi^ NMJs stained for Dlp and HRP. **(m)**. Quantification of Dlp protein relative to bouton area in terminal boutons. N = 13 for w^1118^, 15 for TDP-43^WT^, 16 for TDP-43^G298S^, 6 for TBPH^RNAi^. **(n-y)** Representative images of w^1118^, TDP-43^WT^, TDP-43^G298S^ and TBPH^RNAi^ VNCs stained for Dlp, DNA and TDP-43 **(z)**. Quantification of Dlp granule number in the neuropil. N = 10 for w^1118^, 10 for TDP-43^WT^, 9 for TDP-43^G298S^, 6 for TBPH^RNAi^. Genotypes and stainings, as indicated. Scale bars: **(a)** 10 μm **(n)** 50 μm. **(v)** 22 μm. Significance determined using the Holm Sidak’s multiple comparison test. * = P_value_ < 0.05, ** = P_value_ < 0.01, *** = P_value_ < 0.001. Error bars represent SEM.

These results indicate that TDP-43 proteinopathy causes altered Dlp protein expression, more specifically a reduction at synaptic terminals and aggregate like structures in the VNC neuropil. To investigate whether these alterations are post-transcriptional, as suggested by the RIP and TRAP experiments, we used fluorescence in situ hybridization (RNAScope) to examine *dlp* mRNA localization in the VNC and at the NMJ. Since we were not able to detect a specific signal, possibly due to low levels of expression (data not shown) we performed RT-qPCR experiments using dissected VNCs and larval NMJ preparations and found no significant changes in *dlp* mRNA levels between TDP-43 proteinopathy models and controls (Supplemental Figure 5-1b-c). Although the VNC and NMJ qPCR samples contain multiple cell types, which affects our ability to detect mRNA differences caused by TDP-43 overexpression specifically in motor neurons, these experiments collectively suggest that Dlp expression is likely regulated by TDP-43 post-transcriptionally, possibly through a combination of translation and axonal transport.

### *dlp* is a modifier of TDP-43 dependent locomotor deficits

Next, to test whether Dlp is an effector of TDP-43 proteinopathy, we overexpressed *dlp* using the motor neuron specific driver D42 GAL4 on its own and in the context of TDP-43^WT^ or TDP-43^G298S^ and measured its effect on TDP-43 induced locomotor defects. Using larval turning assays to measure locomotor function we found that although *dlp* overexpression (*dlp*^OE^) on its own resulted in a higher turning time (15.3 +/- 1.01 seconds compared to w^1118^ controls 8.95 +/- 0.46 seconds, P_Value_ = 2E-6), in the context of either TDP-43^WT^ or TDP-43^G298S^ disease models, it caused a lower turning time, indicating a rescue of locomotor dysfunction (11.30 +/- 0.56 seconds compared to 13.40 +/- 0.69 seconds for TDP-43^WT^ alone, P_Value_ = 0.041; 12.50 +/- 0.83 seconds compared to 16.40 +/- 1.16 seconds for TDP-43^G298S^ alone, P_Value_ = 6.4E-3, see Figure 6a). Additionally, although TBPH^RNAi^ knock-down also caused locomotor defects (18.75 +/- 1.69 seconds compared to 8.95 +/- 0.46 seconds for w^1118^, P_value_ = 1.66E-07), *dlp* overexpression in this context is not sufficient to mitigate TBPH^RNAi^ induced locomotor dysfunction (16.13 +/- 1.72 seconds compared to 18.75 +/- 1.69 seconds for TBPH^RNAi^ alone, P_value_ = 0.21, see Figure 6a).

**Figure 6.**
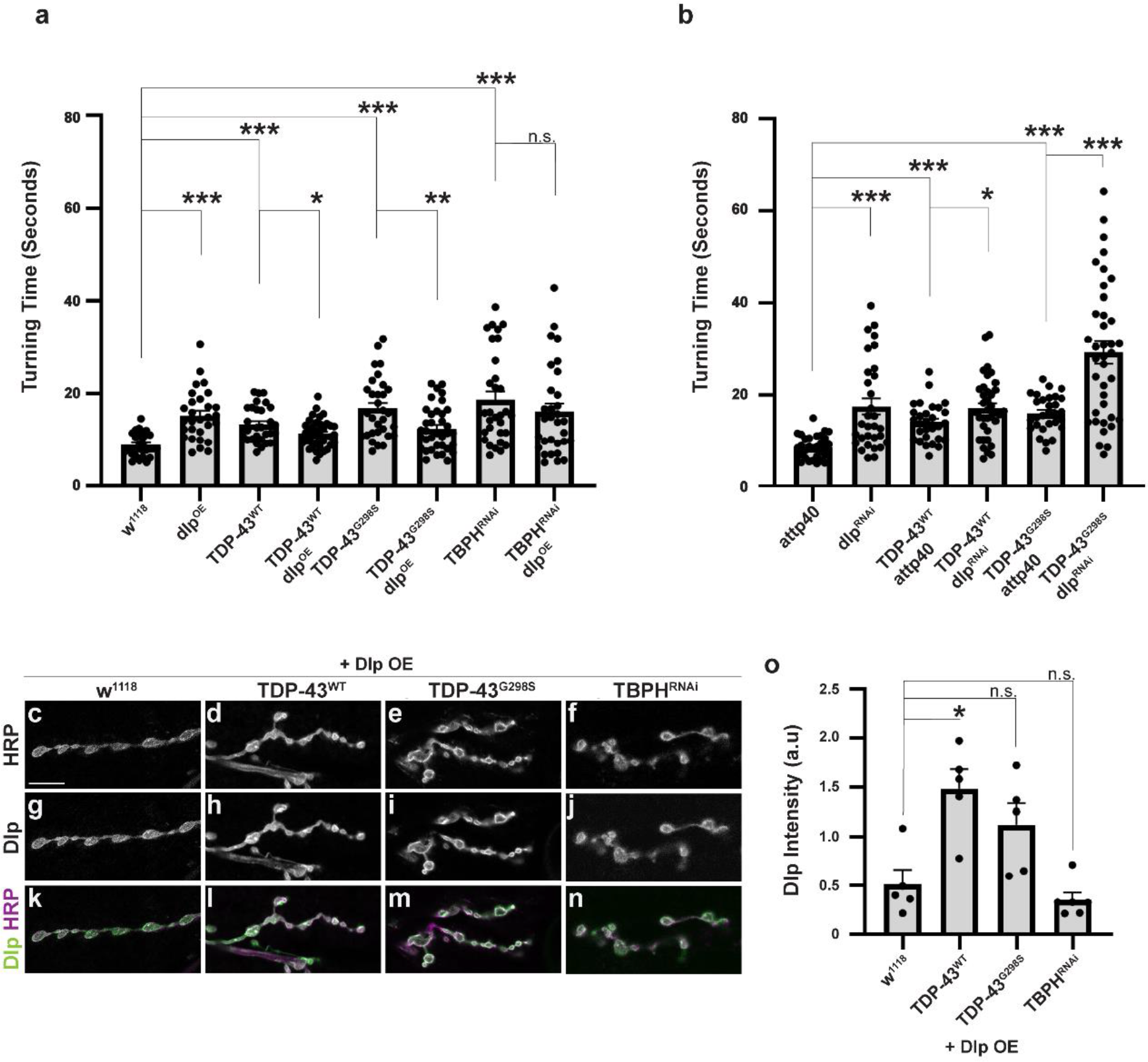
Altered *dlp* mRNA levels modify TDP-43 induced locomotion and lifespan defects. **(a)**. Larval turning times for *dlp* mRNA overexpression in the w^1118^ genetic background (control for OE experiments), TDP-43^WT^, TDP-43^G298S^ and TBPH^RNAi^. N = 29 for w^1118^, 29 for dlp^OE^, 30 for TDP-43^WT^, 37 for TDP-43^WT^ dlp^OE^, 31 for TDP-43^G298S^, 36 for TDP-43^G298S^ dlp^OE^, 32 for TBPH^RNAI^, 31 for TBPH^RNAi^ dlp^OE^. **(b)**. Larval turning times for *dlp* RNAi in the attp40 genetic background (control for RNAi experiments), TDP-43^WT^, and TDP-43^G298S^. N = 30 for attp40, 31 for dlp^RNAi^, 29 for attp40 TDP-43^WT^, 37 for TDP-43^WT^ dlp^RNAi^, 29 for attp40 TDP-43^G298S^, 38 for TDP-43^G298S^ dlp^RNAi^. Significance determined using the MannU Whitney Test or Holm Sidak’s multiple comparison test. * = P_value_ < 0.05, ** = P_value_ < 0.01, *** = P_value_ < 0.001. **(c-n)** Representative images of *dlp*^*OE*^ NMJs in the context of w^1118^, TDP-43^WT^, TDP-43^G298S^ and TBPH^RNAi^ stained for Dlp and HRP. **(o)**. Quantification of Dlp intensity in NMJ terminal boutons. N = 5 for w^1118^ dlp^OE^, 5 for TDP-43^WT^ dlp^OE^, 5 for TDP-43^G298S^ dlp^OE^, 6 for TBPH^RNAi^ dlp^OE^,. Genotypes and stainings, as indicated. Scale bars: **(c)** 10 μm. * = P_value_ < 0.05, ** = P_value_ < 0.01, *** = P_value_ < 0.001. Error bars represent SEM.

Consistent with *dlp*^*OE*^ mitigating TDP-43 induced locomotor defects, *dlp* knockdown by RNA interference (*dlp*^*RNAi*^) enhanced locomotor deficits in both ALS models based on TDP-43 (16.9 +/- 1.21 seconds for TDP-43^WT^ dlp^RNAi^ compared to 14.00 +/- 0.76 seconds for TDP-43^WT^ alone, P_value_ = 0.017; 29.30 +/- 2.56 seconds for TDP-43^G298S^ dlp^RNAi^ compared to 16.10 +/- 0.73 seconds for TDP-43^G298S^ alone, P_value_ = 1.9E-4, see Figure 6b). Interestingly, similar to *dlp*^*OE*^, *dlp*^*RNAi*^ also caused higher larval turning times on its own (17.5 +/- 0.76 seconds compared to 8.64 +/-0.45 seconds for attp40 controls, P_value_ = 1.4E-5, Figure 6b) suggesting that finely tuned levels of Dlp at the NMJ are required for proper locomotor function.

We next asked whether the rescue of TDP-43 induced locomotor dysfunction by *dlp*^*OE*^ could be explained by restoration of Dlp protein levels at the NMJ. To address this, we evaluated Dlp levels within synaptic boutons when *dlp* was overexpressed on its own, in a w^1118^ background (*dlp*^*OE*^) or co-overexpressed with TDP-43^WT^, TDP-43^G298S^ or TBPH^RNAi^ (Figure 6c-n). First we noticed that *dlp*^*OE*^ on its own caused a visible increase (3.88 times higher, P_value_ = 1.654E-6) in Dlp levels at the NMJ compared to w^1118^ controls (Supplemental Figure 6-1a-g) therefore all samples with *dlp*^OE^ were imaged at a lower gain than those without, in order to avoid saturation. While this prevented us from quantitatively comparing samples with and without *dlp*^*OE*^, we were able to evaluate Dlp levels separately, within each set of samples (Figures 5 and 6). These experiments show that Dlp levels in terminal boutons are equal to, or greater than those observed in the *dlp*^*OE*^ alone (TDP-43^WT^, FoldChange = 2.20, P_value_ = 0.0432; TDP-43^G298S^, FoldChange = 1.80, P_value_ = 0.309; TBPH^RNAi^, FoldChange = 0.646, P_value_ = 1.00). Together with our findings that *dlp*^*OE*^ mitigates TDP-43 proteinopathy but not TBPH^RNAi^ induced locomotor defects, these results indicate that Dlp levels at the NMJ are critical in the context of TDP-43 proteinopathy and support a cytoplasmic gain of function effect rather than a nuclear loss of function mechanism. Suprisingly, *dlp*^*OE*^ in the context of TDP-43^WT^ resulted in a statistically significant increase in Dlp levels at the NMJ, suggesting complex regulatory relationship between Dlp and TDP-43 proteinopathy that warrants further investigation.

To further probe the relationship between TDP-43 induced neurodegeneration, Dlp, and Wg/Wnt signaling we used genetic interaction approaches between TDP-43 and the Frizzled 2 (Fz2) receptor, an established Dlp interactor^43^. These experiments showed that while overexpression of wild-type Fz2 had no significant effect on TDP-43 induced locomotor dysfunction (Supplemental Figure 6-1h), RNAi knock-down or overexpression of a dominant negative form of Fz2 caused significant larval lethality in the context of TDP-43 proteinopathy (<10 larvae alive among > 200 total larvae, data not shown). These findings suggest that Wg/Wnt signaling may already be compromised at the NMJ in TDP-43 proteinopathy, albeit the precise mechanism remains to be determined.

### The expression of GPC6, a human Dlp ortholog is altered in patient spinal cords

To validate our findings from *Drosophila* in human patient derived tissues, we evaluated the expression and sub-cellular distribution of the Dlp human ortholog GPC6^80^ in lumbar spinal cord from ALS patients compared to non-neurological controls using immunohistochemistry. These experiments showed that GPC6 appears increased and more granular in ALS spinal cords compared to controls (Figure 7, for patient demographic information see Supplemental Table 7-1). Quantification of staining intensity and granularity showed aggregate-like GPC6 puncta were present in ALS patient motor neurons compared to controls (Fold Change = 7.03, P_value_ = 0.011, see figure 7c). Interestingly, this increased granular appearance of GPC6 in ALS spinal cords resembles the Dlp protein levels and localization observed in the *Drosophila* VNCs overexpressing TDP-43 (Figures 5n-z). Additionally, similar to our findings in the fly models of TDP-43 proteinopathy (Figures 4d, e), soluble vs insoluble fractionations of post-mortem patient tissue revealed that the mRNAs of both *dlp* orthologs *GPC4* and *GPC6* were insolubilized in spinal cords with TDP-43 pathology compared to occipital lobe control tissue from the same patients (N= 4 patients, *GPC4* Log2FC = 2.55, Padj, = 9.62E-17; *GPC6* Log2FC = 1.06, Padj. = 0.0122, Supplementary Table 7-2). Together, these findings show that GPC4/6 alterations in patient tissues resemble those identified in the *Drosophila* models of TDP-43 proteinopathy and suggest that glypican function may also be altered at neuromuscular synapses in ALS, consistent with a recent report of GPC4/6 reduction in SOD1 mice^81^.

**Figure 7.**
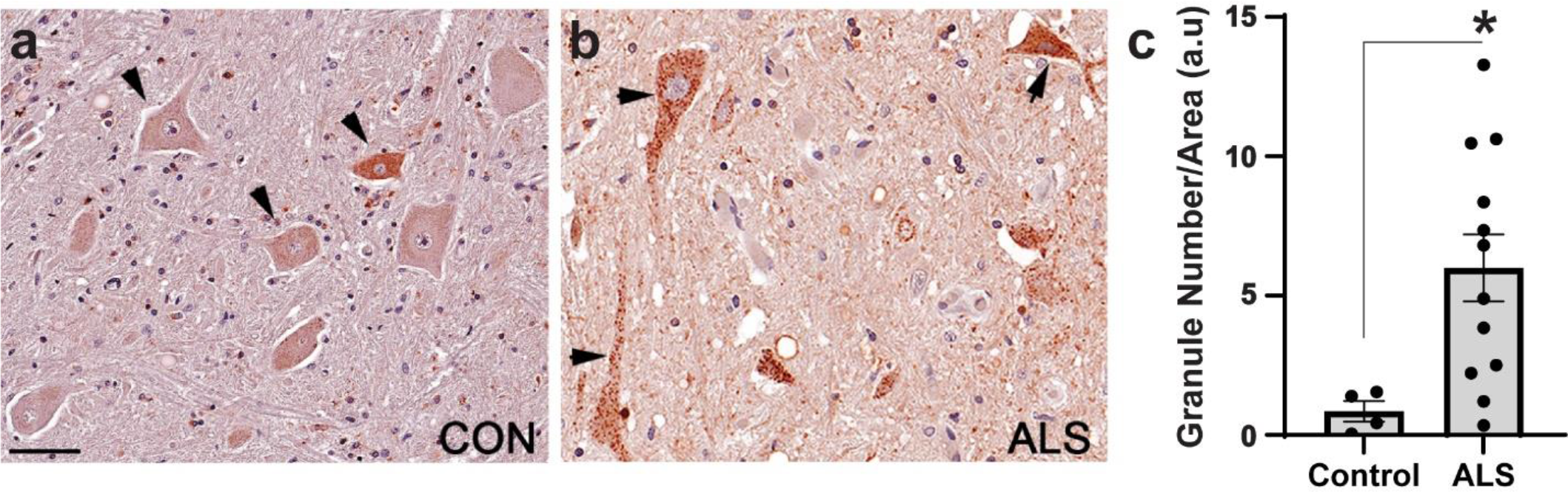
GPC6, a Dlp ortholog exhibits increased expression and altered localization in patient spinal tissues. **(a)**. Representative image of a control spinal cord. N = 4. **(b)**. Representative image of ALS patient spinal cord. N = 12. **(c)** Quantification of GPC6 granule number in post-mortem spinal tissue. Arrowheads indicate motor neurons. Significance determined using the MannU Whitney Test. * = P_value_ < 0.05, Scale bar in **(a)** 40 μm. Error bars represent SEM.

## Discussion

TDP-43 proteinopathy is a hallmark of ALS^13, 82^ and has been observed in several other neurodegenerative diseases^13, 83, 84, 85, 86^, yet its contribution to neurodegeneration remains poorly understood. TDP-43 contributes to multiple RNA processing steps from splicing to translational regulation, providing multiple opportunities for gene expression dysregulation in disease^87, 88^. Furthermore, TDP-43 associates with several cytoplasmic complexes including translational machinery^89, 90, 91, 92^ and stress granules^19, 36, 93, 94^, suggesting a role for TDP-43 in translation inhibition, as predicted by the ribostasis hypothesis^42^. Several specific mRNA targets of TDP-43 mediated translational inhibition have been reported^24, 39, 40, 95^. Among these, *futsch/MAP1B* and *Hsc70-4/HSPA8* modify disease phenotypes highlighting their contribution to disease^24, 39^. Recently, puromycin incorporation experiments in SH-SY5Y neuroblastoma cells showed that increased cytoplasmic TDP-43 reduces global translation through interactions with RACK1 on polyribosomes^90^. In contrast, polysome fractionation experiments in a human cell model showed that ALS associated mutant TDP-43^A315T^ acts as a positive regulator of translation for a subset of specific mRNAs (*e*.*g*., Camta1, Mig12, and Dennd4A)^41^. Taken together these findings highlight a complex role for TDP-43 in translational regulation that can act both as a translational activator and inhibitor.

Here, using a *Drosophila* model of TDP-43 proteinopathy that recapitulates key features of the human disease including locomotor dysfunction and reduced lifespan^45, 46^ we identified TDP-43 dependent changes in the motor neuron translatome. These experiments identified both pathways previously associated with ALS (*e*.*g*., translation, mitochondrial function^96, 97^) and novel targets of TDP-43 mediated translational inhibition including the Wg/Wnt signaling regulator *dlp* and the glutathione metabolism pathway. Although glutathione dysregulation has been shown to be deficient both in patient tissue^98^ and patient derived neuronal cultures^99^, it has not previously been identified as a direct consequence of TDP-43 pathology. Surprisingly, genes related to cytoplasmic translation are enriched in the ALS motor neuron translatomes relative to the RpL10 controls, possibly reflecting a compensatory mechanism whereby degenerating neurons upregulate global translation in response to increased cellular stress or to mitigate cytoplasmic TDP-43’s inhibitory effect on protein synthesis^90^.

In addition to identifying motor neuron specific alterations to the translatome induced by TDP-43 proteinopathy, we identified *dlp* as a novel target of TDP-43. *dlp* is translated into a GPI anchored glypican that interacts with the Wg/Wnt pathway receptor Fz2, serving as both a co-factor and competitive inhibitor for substrate binding^43^. RNAi knockdown or overexpression of a dominant negative form of Fz2 resulted in lethality prior to the third instar stage in the context of both TDP-43^WT^ and TDP-43^G298S^ disease models suggesting that the Wg/Wnt signaling pathway may be compromised by TDP-43 proteinopathy as suggested by expression studies in patient tissues^76, 77^. Although overexpression of wild-type Fz2 did not mitigate TDP-43 dependent locomotor defects as would be expected based on the lethality caused by Fz2 loss of function, it is possible that cell autonomous (*i*.*e*., motor neuron) and non cell autonomous (*i*.*e*., glia, muscle) aspects of Wg/Wnt signaling^74^ may be at play and confound our genetic interaction experiments. Alternatively, Wg/Wnt signaling and TDP-43 proteinopathy act in parallel pathways. Indeed, Dlp could be an effector of TDP-43 proteinopathy through interactions with multiple pathways including the hedgehog^100^ and hippo^101^ pathways that Dlp function has also been implicated in.

We show that although TDP-43^WT^ and TDP-43^G298S^ both associate with *dlp* mRNA, and correlate with *dlp* depletion from ribosomes, the severity of some phenotypic alterations was variant dependent. While overexpression of *dlp* mRNA was sufficient to rescue locomotor dysfunction and Dlp protein levels were altered in the context of both TDP-43 variants, *dlp* mRNA was insolubilized only in the TDP-43^G298S^ model. Additionally, different pathways were shown to have altered translation between the TDP-43^WT^ and TDP-43^G298S^ models. Differential results have previously been observed between the TDP-43^WT^ and TDP-43^G298S^ models^39^, which are likely the result of increased TDP-43^G298S^ stability^102^ and granule viscosity relative to TDP-43^WT,103^.

In both ALS models, analyses of the whole VNC, which contains motor neuron soma and neurites, show that Dlp forms distinct puncta whereas staining at the NMJ revealed a significant reduction in Dlp protein. These findings suggest that TDP-43 dependent translational inhibition may be localized to axons and NMJs, which can explain the relatively low magnitude depletion of *dlp* mRNA from ribosomes using TRAP, an approach that reflects translational changes in the entire motor neuron rather than specific compartments. It is possible that the synapse specific alteration in Dlp levels could also result from transport deficits, trapping NMJ-bound *dlp* mRNA/Dlp protein in the soma or neurites. Although our qPCR data show no change in steady state *dlp* mRNA levels, we cannot eliminate the possibility of mRNA and/or protein transport defects leading to the accumulation of Dlp in the VNC and reduction at the NMJ.

Interestingly, TBPH^RNAi^ was sufficient to deplete Dlp from the NMJ, but not induce significant neuropil puncta, suggesting that the two phenotypes can occur independently of each other and might result from different mechanisms. Loss of nuclear function versus cytoplasmic gain of function remains an open question, with both mechanisms likely contributing to TDP-43 proteinopathy^32, 104^. Dlp depletion at the NMJ in the context of TBPH^RNAi^ suggests that this phenotype is the result of nuclear TDP-43 function whereas the neuropil puncta are likely the result of toxic gain of function as they are significant only in the context of TDP-43 proteinopathy and not in the context of endogenous TDP-43 loss of function. Furthermore, the fact that restoring Dlp expression at the NMJ is not sufficient to mitigate TBPH^RNAi^ induced locomotor defects while it improves TDP-43^WT^ and TDP-43^G298S^ dependent phenotypes suggests that Dlp levels at the NMJ are critical in the context of TDP-43 proteinopathy and that other factors may be at play in the loss of function caused by TBPH^RNAi^ knock-down. Collectively, our findings suggest a model in which the mislocalization of Dlp found in flies and patient spinal cords results from a combination of translation inhibition and transport defects as previously reported for Futsch/MAP1b^24^ (see Figure 8 for model). Lastly, although more experiments are needed to determine the nature and composition of Dlp puncta we found in the VNC neuropil, we speculate that they may reflect the accumulation of endomembrane compartments that were previously shown to be altered in the context of TDP-43 proteinopathy^105^.

**Figure 8.**
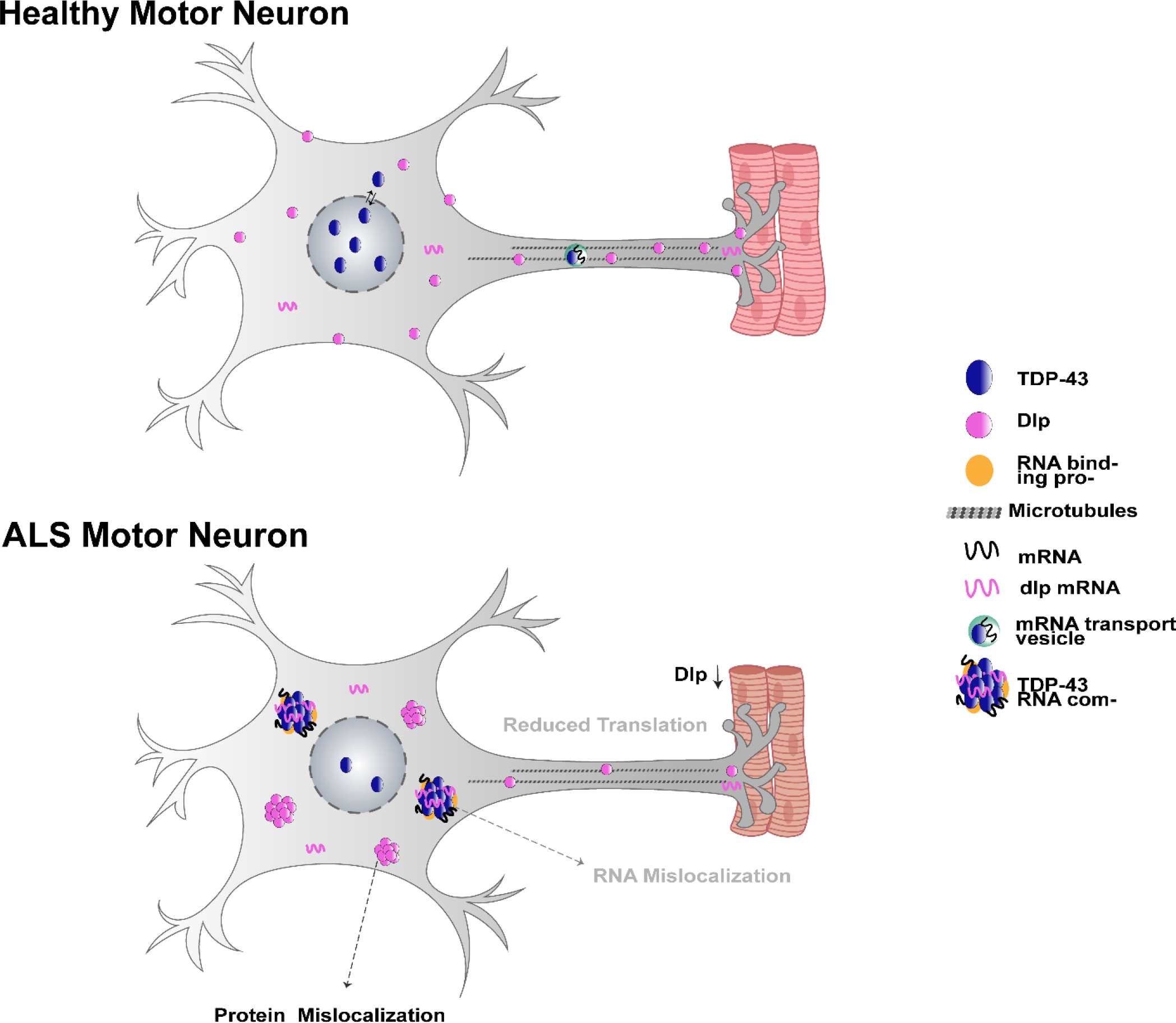
Dlp is altered in the context of TDP-43 proteinopathy. Dlp mRNA is enriched in TDP-43 complexes, depleted from ribosomes and sequestered in insoluble/urea complexes. Taken together, these findings and our observations of Dlp protein being reduced at the NMJ and accumulating in puncta within cell bodies and neuropils suggest multiple cellular defects including local translation inhibition at synapses, axonal trafficking and endomembrane trafficking deficits.

Our findings that GPC6, a Dlp ortholog, is increased in ALS spinal cord motor neurons, insolubilzed in patient spinal tissues, and has puncta resembling the Dlp granules observed in the *Drosophila* VNC, further highlighs the power of the fly models to predict pathological changes in disease. Interestingly, the formation of heparan sulfate proteoglycan containing puncta has previously been observed in other neurodegenerative disease including Alzheimer’s, Parkinson’s and SOD1 mediated ALS (reviewed in^106^) however this has not been previously reported in the context of TDP-43 proteinopathy, nor has a mechanism been proposed. More recently, genetic knockdown of heparan sulfate-modifying enzyme *hse-5* in C. elegans mitigated TDP-43 induced deficits in synaptic transmission^107^, further suggesting a link between TDP-43 pathology, neurodegeneration and heparan sulfate proteoglycans such as Dlp/GPC6.

In summary, we report several novel translational consequences of TDP-43 proteinopathy including on the spliceosome, metabolic pathways and the translational machinery itself. We also identify the glypican *dlp* as a target of TDP-43 proteinopathy, which highlights multiple therapeutic strategies including restoration of synaptic specific protein expression, intracellular transport as well as Wg/Wnt and other Dlp linked signaling pathways in ALS.

## Materials and Methods

### Drosophila genetics

*w*^*1118*^; *UAS-TDP-43*^*WT*^*-YFP* and *w*^*1118*^; *UAS-TDP-43*^*G298S*^*-YFP* were previously described^45, 46^. *Drosophila* harboring UAS-RpL10 GFP were obtained from Herman Dierick^47^. *w*^*1118*^; *UAS-TDP-43*^*WT*^ *UAS-RpL10-GFP/CyO* and *w*^*1118*^; *UAS-RpL10-GFP/CyO; UAS-TDP-43*^*G298S*^*/TM6B* were generated with *w*^*1118*^; *UAS-TDP-43*^*WT* 108^ and *w*^*1118*^; *UAS-TDP-43*^*G298S* 109^ using standard genetic approaches. *w*^*1118*^; *UAS-TBPH*^*RNAi*^ was generated by recombining *y[1] v[1]; P{y[+t7*.*7] v[+t1*.*8]=TRiP*.*HMS01846}attP40* (Bloomington Stock 38377) and *y[1] v[1]; P{y[+t7*.*7] v[+t1*.*8]=TRiP*.*HMS01848}attP40* (Bloomington Stock 38379). These lines were crossed with the D42 GAL4 driver^110^ to achieve motor neuron specific expression. For *dlp* overexpression we used *w*^*1118*^; *P{w[+mC]=UAS-dlp*.*WT}3* (Bloomington Stock #9160) and for RNAi knock-down we used *y[1] v[1]; P{y[+t7*.*7] v[+t1*.*8]=TRiP*.*GLC01658}attP40* (Bloomington Stock #50540). As a control for the RNAi experiments we used *y[1] v[1]; P{y[+t7*.*7]=CaryP}attP40* (Bloomington Stock #36304). CRISPR TDP-43 models were previously described^78^ and provided by David Morton. For Fz2 overexpression we used *w[*]; P{w[+mC]=UAS-fz2-2}16/TM6B, Tb[+] (*Bloomington Stock #41794. For Fz2 knockdown we used *y[1] sc[*] v[1] sev[21];P{y[+t7*.*7]v[+t1*.*8]= TRiP*.*HMS05675}attP40* (Bloomington Stock #67863). Expression of dominant negative Fz2 was achieved with *w[*]; P{w[+mC]=UAS-fz2(ECD)-GPI}3* (Bloomington Stock #44221).

### RNA immunoprecipitations (RIP)

100 third instar larvae expressing TDP-43^WT^-YFP or TDP-43^G298S^-YFP were collected and flash frozen in liquid nitrogen. Frozen larvae were homogenized in lysis buffer (100 mM HEPES Buffer pH 8.0, 1% Triton X-100, 200 mM NaCl, 30 mM EDTA, 350 mM Sucrose, 10% Glycerol, 1 mg/mL Heparin, 1 mM DTT, protease inhibitors (Millipore Sigma 11873580001) and RNAsin Plus 400 units/ml (Fischer Scientific PRN2615). Lysates were centrifuged at 10,000 x g for 10 minutes and then pre-cleared with magnetic beads (Dynabeads Protein A, Thermofisher Scientific 10001D) for 1 hour at 4°C. Magnetic beads bound to chicken anti-GFP antibody (Life tech A-11122) were added to the supernatant followed by rotation at 4 °C for 2 hours. Next, beads were washed 3 times with a sugar-rich buffer (100 mM HEPES Buffer pH 8.0, 1% Triton X-100, 200 mM NaCl, 30 mM EDTA, 350 mM Sucrose, 10%, Glycerol, 1 mg/mL Heparin, 1 mM DTT, RNAsin Plus 400 u/ml) and 2 times with a low-density buffer (100 mM HEPES Buffer pH 8.0, 1% Triton X-100, 200 mM NaCl, 30 mM EDTA, 1 mg/mL Heparin, 1 mM DTT, RNAsin Plus 400 units/ml (Fischer Scientific PRN2615) followed by resuspension in Qiagen RLT buffer (Qiagen 79216) with 1% 2-mercaptoethanol (Sigma Aldrich M6250).

### Tagged ribosomes affinity purifications (TRAP)

RNA associated with ribosomes was immunoprecipitated using a protocol similar to RIP (see above) except that the genotypes were *w*^*1118*^; *UAS-RpL10-GFP/CyO, w*^*1118*^; *UAS-TDP-43*^*WT*^ *UAS-RpL10-GFP/CyO* and *w*^*1118*^; *UAS-RpL10-GFP/CyO; UAS-TDP-43*^*G298S*^*/TM6B Hu Tb* and the buffers for the immunoprecipitation of RpL10-GFP also included 100 μg/mL cycloheximide.

### RNA Isolation

RNA was isolated from the ventral cord lysates, whole larvae inputs, or immunoprecipitations using a Qiagen RNeasy Mini Kit (Qiagen 74104) with quality and quantity determined by a nanodrop spectrophotometer. For fractionation experiments and dissected ventral nerve cords (VNCs) or neuromuscular junctions (NMJ), RNA was isolated using Trizol (Thermofisher Scientific 15596026).

### RNA Seq

Isolated RNA was prepared for RNA sequencing using the SMART-seq v4 Ultra Low Input RNA kit (Takara Bio USA 634888). The Nextera XT DNA Library Prep Kit (Illumina FC-131-1024) was used to tag, clean, and pool the samples with quality assayed by an Agilent 2100 Bioanalyzer. The cDNA libraries were sequenced by the Beijing Genomics Institute using an Illumina HiSeq 4000. Human fractionation RNA-seq were performed at the Translational Genomics Institute (TGen).

### RNA-seq Analysis

Reads were aligned to the Flybase FB2018_05 genome using MAFFT^111^ with differential expression analysis conducted through Deseq2^112^. In RIP experiments, 9812 and 9954 genes were detected in TDP-43^WT^ and TDP-43^G298S^ complexes, respectively. In TRAP experiments, 9711, 9347, and 9669 genes were detected in RpL10 GFP complexes in the control, TDP-43^WT^, and TDP-43^G298S^ models, respectively. To identify the “normal” motor neuron translatome we calculated the Log2FoldChange between RpL10 IP and RpL10 VNCs. Next, to identify alterations resulting caused by TDP-43 proteinopathy to the motor neuron translatomes, we subtracted the Log2FoldChange between RpL10 IP and RpL10 VNCs from the Log2FoldChange between TDP-43^WT^ or ^G298S^ RpL10 IP and TDP-43^WT^ or ^G298S^ RpL10 IP VNCs. For comparison, we also normalized RpL10 IPs to RpL10 whole larvae inputs (see Supplemental Figure 1). Gene ontology (GO) analyses were conducted using David 6.8^49, 50^. Human spinal cord transcriptomic data underwent Deseq2 analysis from counts provided by target ALS, detecting 33843 genes. For the human fractionation RNA-seq data, reads were aligned to the NCBI GRCh38 genome with gencode 29 genome using STAR^113^ then underwent Deseq2 differential expression analysis detecting 24090 genes. GO plot 1.0.2^114^, STRING 11.0^73^, and the R package “LPS” version 1.0.10 were used to generate bio-informatic figures.

### Fractionations

Fractionations were conducted as previously described^39^. In brief, 25 third instar larvae were homogenized in Trizol (Thermofisher Scientific 15596026). Lysates were then centrifuged at 25,000 x g for 30 minutes. The supernatant became the soluble fraction and the pellet was solubilized in urea buffer (30 mM Tris, 7 M Urea, 2 M Thiourea, 4% CHAPS, 1X Protease Inhibitor Cocktail (Millipore Sigma 11873580001), 0.5 mM PMSF, RNAsin Plus 400 units/ml (Fischer Scientific PRN2615), pH = 8.5) to generate the urea/insoluble fraction. For human fractionation samples, post-mortem tissue (four spinal cord and two cerebellum samples as controls) was homogenized and subjected to the fractionation protocol described above.

### Immunofluorescence (*Drosophila* tissues)

Third instar ventral nerve cords were dissected as previously described^24^. Primary antibodies/stains used were 1:5 anti-Dlp (antibody 13G8, developed by Phil Beachy, obtained from the Developmental Studies Hybridoma Bank, created by the NICHD of the NIH and maintained at The University of Iowa, Department of Biology, Iowa City, IA 52242) and 1:200 anti-GFP (Rockland 600-102-215). DNA was visualized using Hoechst (1:10,000, Invitrogen H3570). Goat anti-mouse Alexa 568 (1:500 Thermofisher Scientific A-11004) was used as secondary antibody. Samples were imaged using a Zeiss LSM 880 inverted confocal microscope with a 40X oil lens. The number and cumulative area of Dlp puncta was quantified via manual counting and outlining of blinded VNC images. Third instar larval NMJs were dissected as previously described^24^. Primary antibodies used were 1:5 anti-Dlp, 1:100 anti-HRP Alexa 647 (Jackson Immuno-research 123-605-021). Goat anti-mouse Alexa Fluor 568 (1:500 Thermofisher Scientific A-11004) was used as secondary antibody. Samples were imaged using a Zeiss LSM 880 NLO upright confocal microscope with a 40X oil lens. Dlp intensity per area was quantified with NIH imageJ v1.52p within four terminal boutons defined by HRP staining. All images were blinded prior to being analyzed.

### Immunohistochemistry (human post-mortem spinal cords)

Post-mortem spinal cord tissue sections were obtained from the Barrow Neurological Institute ALS Tissue Bank and Target ALS Postmortem Tissue Core. All tissue samples were collected after informed consent from the subjects or by the subjects’ next of kin, complying with all relevant ethical regulations and approved by the appropriate Institutional Review Boards. Clinical neuropathological diagnoses were made by board certified neuropathologists according to consensus criteria. Subject demographics are listed in Figure 7, Supplementary Table 1. A total of 12 ALS and 4 controls (4 non-neurologic disease controls) were used in this study. Tissue sections were processed and imaged as previously described^115^. All ALS cases contained TDP-43 pathology while the non-neurologic disease controls lack TDP-43 pathology. All sections were stained for GPC6 protein using rabbit polyclonal primary antibody at 1:300 (Bioss bs-2177R). A biotinylated goat anti-rabbit IgG (H+L) was used as secondary antibody (Vector Laboratories, BA-1000, 1:200). Images were blinded, then motor neurons were outlined and inverted in NIH ImageJ, which was also used to quantify GPC6 protein intensity relative to background. CellProfiler 3.1.9^116^, was used to quantify GPC6 puncta, defined between 3 to 13 pixels in diameter.

### qRT-PCR

All cDNA was synthesized using Fisher First Strand cDNA synthesis reagents (Thermofisher Scientific K1641). qPCR reactions were conducted in three biological replicates using Taqman Fast Advanced Master Mix (Thermofisher Scientific 4444556) and conducted on either an ABI 7100 or Analytik Jena 844-00504-4 qTOWER qPCR machine. *dlp* mRNA was quantified using Taqman assay (Thermofisher Scientific Dm01798597_m1) using *GPDH* (Thermofisher Scientific Dm01841185_m1) for normalization. Fold change was calculated using the standard ΔΔCT method^117^.

### Locomotor assays

Third instar wandering larvae were placed on a grape juice agar plate. After being given 1 minute to acclimate, the larvae were turned on the ventral side and timed until they uprighted themselves and began the first forward motion^24^. Data points outside of the median turning time +/- 1.5IQR within each genotype were identified as outliers and excluded from statistical analyses.

### Fluorescent In Situ Hybridization (FISH)

FISH was performed using RNAscope (ACD Bio). A custom probe for dm-dlp and a control probe dm-Gapdh1 were used to detect *dlp* and *Gapdh1* in dissected VNC and NMJs as previously described^118^. Signals were detected using a Zeiss LSM 880 inverted confocal microscope with a 40X oil lens.

### Statistical Analyses

For RNA seq analyses, significance was determined using the P adjusted value calculation embedded within Deseq2. All other statistical analyses were performed using GraphPad Prism 7.0 using two-tailed analyses. Normality was evaluated using the Kolmogorow-Smirnov test. All data shown are mean ± standard error of the mean.

### Data Availability

*Drosophila* RNAseq data is submitted at the NCBI GEO database (GSE156222). Human RNAseq data is in the process of being submitted at the NCBI dbGaP database.

### Code Availability

Code is available upon request. Trimming, alignment, and differential analysis was conduced by Eric Alsop and figure generation was conducted by Erik Lehmkuhl.

## Acknowledgements

We thank members of the Zarnescu lab, including Samantha Macklin. Taylor Wingfield, Rebekah Keating Godfrey, Zachary Hammer. Ernesto Manzo, Matthew Scandura, Josh Paree, Alyssa Coyne, and Ben Zaepfel for help with dissections and comments on the manuscript. We also thank Herman Dierick, Takeshi Iwatsubo, Paul Taylor and David Morton for providing Drosophila strains. We acknowledge the Bloomington Drosophila Stock Center and the University of Iowa Developmental Studies Hybridoma Bank for supplying reagents. We thank the Barrow Neurological Institute ALS Tissue Bank and Target ALS Postmortem Tissue Core for access to tissues and slides, and all patients for participating in this study. We also thank the New York Genome Center ALS Consortium for providing access to the RNA-seq database. Funding was provided by National Institutes of Health NIH NS091299, Sandra Harsha Estate (to DCZ), the ARCS foundation and NIH T32-GM008659 (to EML), the Undergraduate Biological Research Program (to ADB, NM, HB and MEM), the Beckman Scholars Program (to RJE), the UArizona Graduate College Award (to CK), a Muscular Dystrophy Association B2I award (to AJ), Target ALS Foundation (to RB) and donations on behalf of ALS patients (to KJ).

## Figure legends

**Supplemental Figure 5-1.**
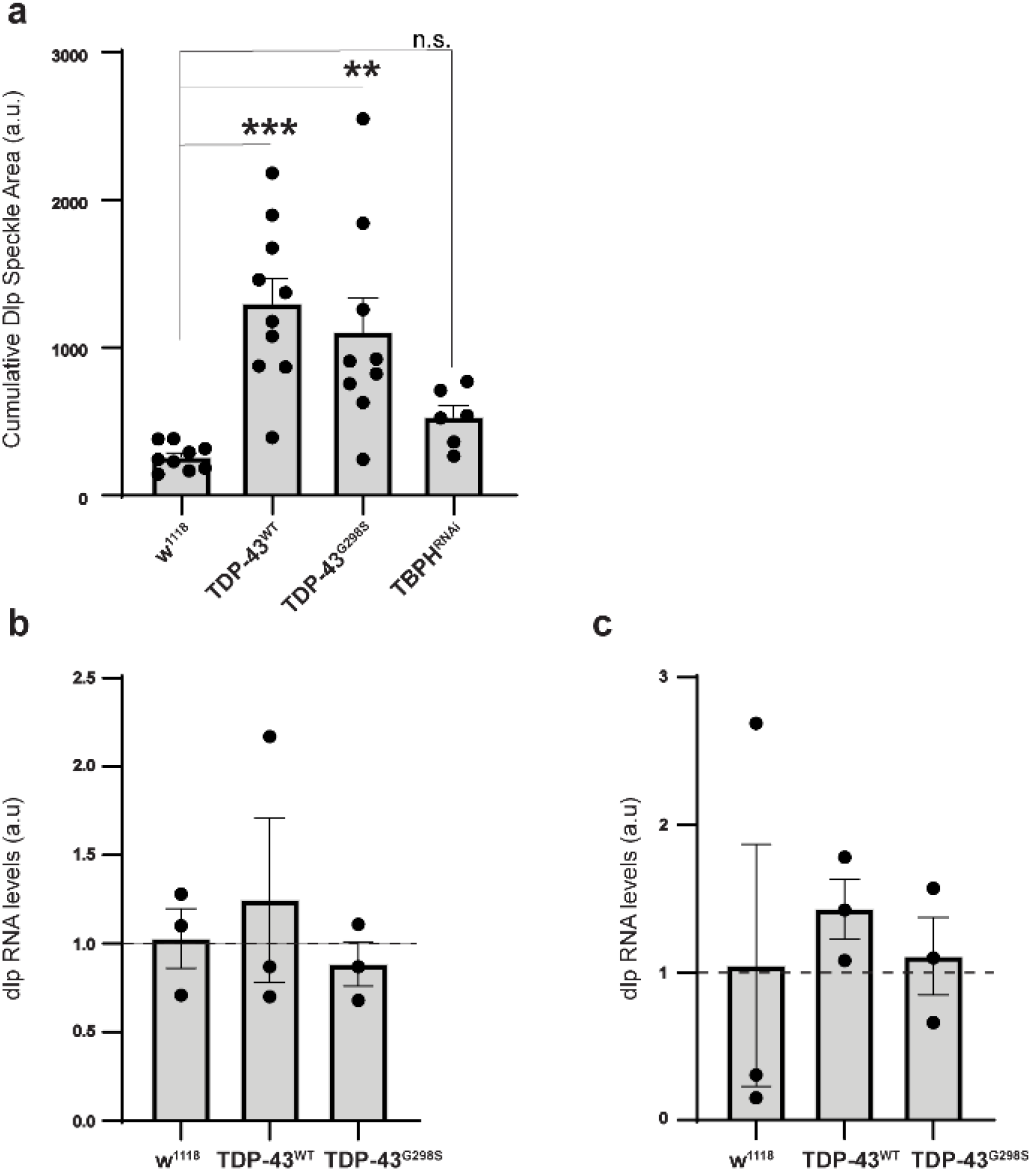
Alterations to *dlp* RNA levels were not detected in either the NMJ or the whole VNC. **(a)**. Cumulative speckle area of Dlp granules in the VNC neuropil from figure 5. N = 9 for w^1118^, 10 for TDP-43^WT^, 9 for TDP-43^G298S^, 6 for TBPH^RNAi^. **(b)**. Quantification of *dlp* mRNA relative to *GPDH* in dissected VNCs. N = 3. (**c)**. Quantification of *dlp* mRNA relative to *GPDH* in dissected NMJs. N = 3. Significance determined through Holm Sidak’s multiple comparison test. * = Pvalue < 0.05, ** = Pvalue < 0.01, *** = Pvalue < 0.001. Error bars represent SEM.

**Supplemental Figure 6-1.**
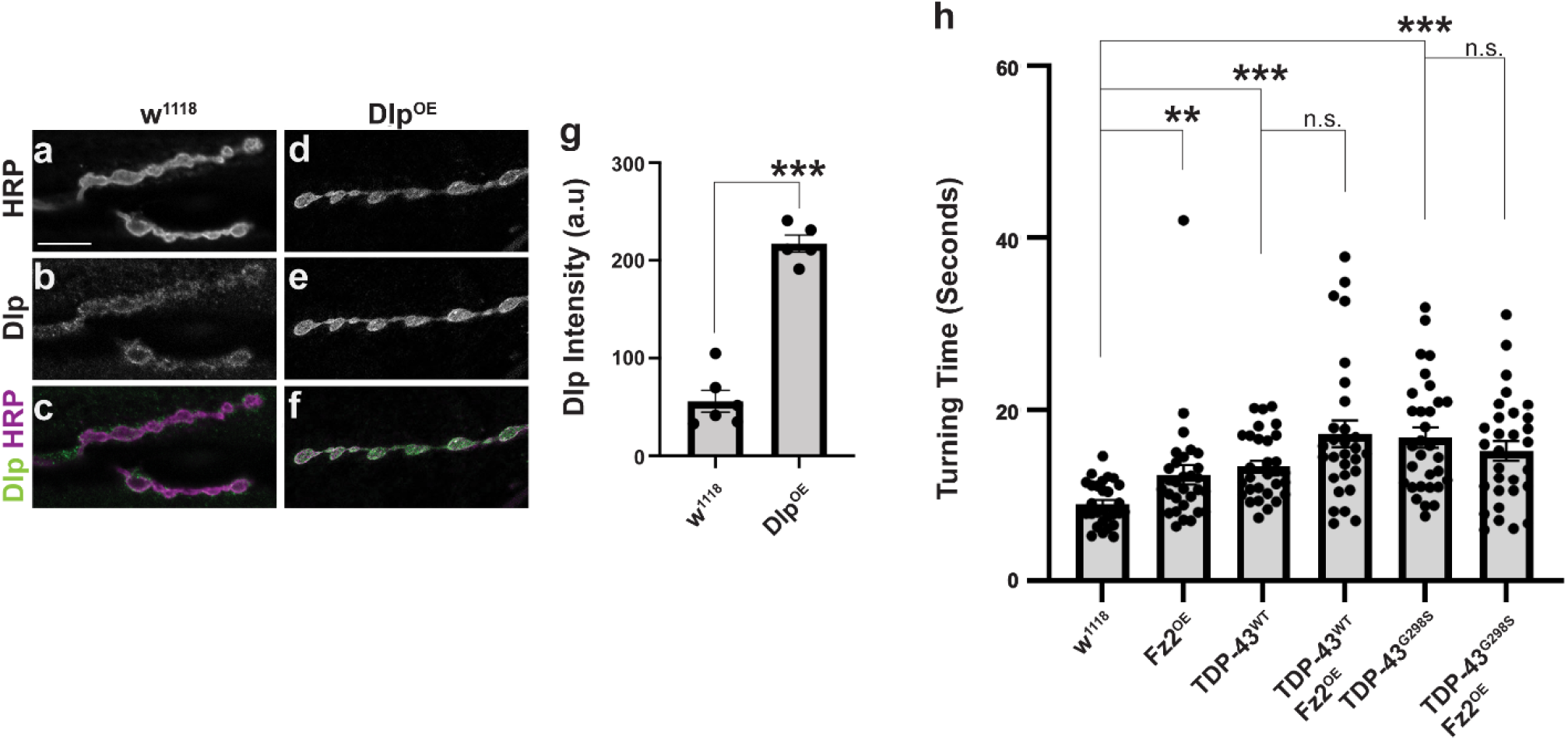
Cumulative Dlp speckle area in the context of TBPH-RNAi. **(A). (a-f)** Representative images of w^1118^ and Dlp^OE^ NMJs stained for Dlp and HRP. **(g)**. Quantification of Dlp granule number in the neuropil. N = (w^1118^ = 6, dlp^OE^ = 5). Genotypes and stainings, as indicated. Scale bars: **(a)** – 10 um. Significance determined through student’s T Test. (t_value_ = 10.96, degrees of freedom = 9). * = Pvalue < 0.05, ** = Pvalue < 0.01, *** = Pvalue < 0.001 **(h)** Larval turning times for *Fz2* mRNA overexpression in the w^1118^ genetic background (control for OE experiments), TDP-43^WT^, and TDP-43^G298S^. N = 29 for w^1118^, 30 for Fz2^OE^, 30 for TDP-43^WT^, 29 for TDP-43^WT^ Fz2^OE^, 31 for TDP-43^G298S^, 30 for TDP-43^G298S^ Fz2^OE^. Significance determined using the MannU Whitney Test or Holm Sidak’s multiple comparison test. * = Pvalue < 0.05, ** = Pvalue < 0.01, *** = Pvalue < 0.001. Error bars represent SEM.

**Figure 4 – Supplemental Table 1.** *dlp* enrichment with TDP-43 (RIP) and depletion from the ribosome in ALS models relative to the RPL10 control (TRAP).

|  | RIP TDP-43WT |  | RIP TDP-43--G298S |  | TRAP TDP-43-WT |  | TRAP TDP-43-G298S |  |
| --- | --- | --- | --- | --- | --- | --- | --- | --- |
| Gene | Log2FC | Padj. | Log2FC | Padj. | Log2FC | Padj. | Log2FC | Padj. |
| *dlp* | 3.62 | 8.32E-57 | 4.63 | 2.47E-57 | -0.394 | 0.0274 | -0.213 | 0.0121 |

**Figure 7 – Supplemental Table 2.**
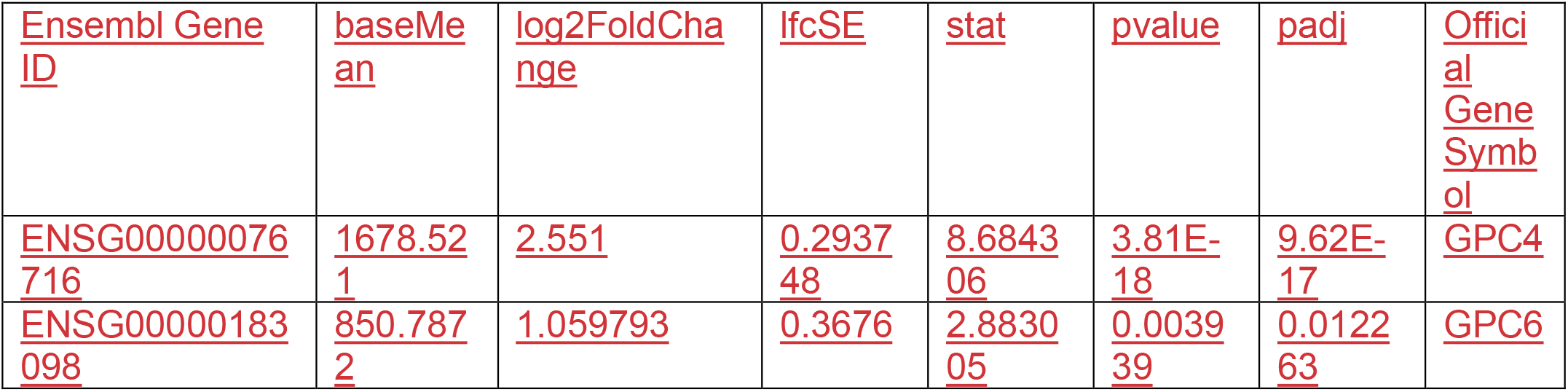
*GPC4/GPC6* differential expression analysis comparing the urea fractionation of patient spinal tissue containing TDP-43 pathology to the urea fractionation of control tissue.

